# Alzheimer’s Disease risk modifier genes do not impact tau aggregate uptake, seeding or maintenance in cell models

**DOI:** 10.1101/809137

**Authors:** Sourav Kolay, Marc I. Diamond

## Abstract

Alzheimer’s disease (AD) afflicts millions of people worldwide, and is caused by accumulated amyloid beta and tau pathology. Progression of tau pathology in AD may utilize prion mechanisms of propagation in which pathological tau aggregates released from one cell are taken up by neighboring or connected cells and act as templates for their own replication, a process termed “seeding.” In cultured cells we have modeled various aspects of pathological tau propagation, including uptake of aggregates, induced (naked) seeding by exogenous aggregates, seeding caused by Lipofectamine-mediated delivery to the cell interior, and chronic maintenance of aggregates in cells through mother-to-daughter transmission. The factors that regulate these processes are not well understood, and we hypothesized that AD risk modifier genes might play a role. We identified 22 genes strongly linked to AD via meta-analysis of genome-wide association studies (GWAS). We used CRISPR/Cas-9 to individually knock out each in gene in HEK293T cells, and verified disruption using genomic sequencing. We then tested the effect of gene knockout in tau aggregate uptake, naked and Lipofectamine-mediated seeding, and aggregate maintenance in cultured cell lines. GWAS gene knockouts had no effect on these models of tau pathology. With obvious caveats due to the model systems used, these results imply that these 22 AD risk modifier genes do not directly modulate tau uptake, seeding, or aggregate maintenance.

## INTRODUCTION

Tauopathies are neurodegenerative diseases characterized accumulation of tau protein in ordered assemblies. Tauopathy progresses according to predictable patterns in patients(1) and has been proposed to involve brain networks(2,3). Our initial studies described the diversity of self-propagating fibrillar conformations *in vitro*(4), and the ability of tau aggregates to propagate pathology from the outside to the inside of a cell, and between cells(5). Concurrent work from the Tolnay group demonstrated that inoculation of mouse brain with tau aggregates induced local pathology in a transgenic mouse model(6). This led us initially to propose that tau had properties similar to the prion protein, PrP(7). In subsequent work, we propagated distinct tau strains in cultured cells that we used to create transmissible tauopathy in mouse models, with faithful, inter-animal propagation of defined pathology(7). This was the first evidence that an infectious form of tau created *in vitro*, would faithfully transmit unique conformations between animals, and we henceforth referred to tau as a prion. This idea remains controversial(8,9). Nonetheless, similar results from multiple groups (10-12) and the effectiveness of immunotherapies against tau in mouse models(13) have now led to a general recognition of the idea that transcellular propagation of pathology could underlie pathogenesis of tauopathies and other amyloidoses. The precise mechanisms are unknown.

Genome-wide association studies (GWAS) have been used to identify risk modifying genes in AD(14). Thousands of individuals with AD have been evaluated, and a relatively small number of genes have been consistently identified (15,16). We hypothesized that one explanation for increased AD risk could be modulation of transcellular propagation of tau pathology. This is very labor-intensive to study in cultured neurons or animal models. Consequently, we have developed cell-based assays to study various steps in tau propagation: uptake(17),(18) conversion of intracellular tau to an aggregated state(17), (19) and maintenance of intracellular aggregates through mother-daughter transmission in dividing cells (7,20). We therefore tested the impact of AD GWAS genes through systematic genetic knockout via CRISPR/Cas-9.

## MATERIALS AND METHODS

### Generation of CRISPR/Cas9 knockout cells and lentiviral transduction

Two human gRNA sequences per gene were selected from the optimized GeCKO version 2(26) or Brunello libraries (27). DNA oligonucleotides were synthesized (IDT), and cloned into the lentiCRISPR v2 vector (26) for lentivirus production. Lentivirus was created as described previously (28). For transduction, a 1:30 dilution of virus suspension was added to the cells. After 24h infected cells were treated with 1μg/ml puromycin (Life Technologies) and cultured for 2 days, followed by passaging 1:5 and a second round of virus and puromycin application. The cells were cultured at least 10 days after the first lentiviral transduction before using them for experiments.

### Confirmation of gene editing by TIDE

Two gRNAs for each gene were used to produce knockout cell lines for analysis by TIDE to confirm the presence of indels in predicted DNA regions as established by Brinkman *et al*. Genomic DNA was extracted (Qiagen DNeasy Blood & Tissue Kit). DNA concentration was determined by spectrophotometer (DeNovix DS-11 FX+). PCR primers were designed around the region of expected CRISPR/Cas 9 cut site according to the protocol for TIDE. PCR was performed using 100ng of genomic DNA with 2x TaqPro Red Complete Polymerase (Denville Scientific). PCR conditions were at 95 °C, then 30X (15s at 95°C, 15s at 60°C, 1min at 72°C) and 10 min at 72°C. The PCR product was run on a 1% agarose gel to verify the product size and gel-extracted using the QIAquick gel extraction kit (Qiagen). Purified PCR samples were Sanger sequenced at the sequencing core facility at UT Southwestern Medical Center. Sequencing files were used for TIDE. Analysis was performed according to the software instructions. The presence of aberrant sequence signal, *R*^2^ value and the knockout efficiency were considered to evaluate the results(22). One gRNA was selected for each gene based on its gene knockout efficacy (Table 1).

**Table 1:**
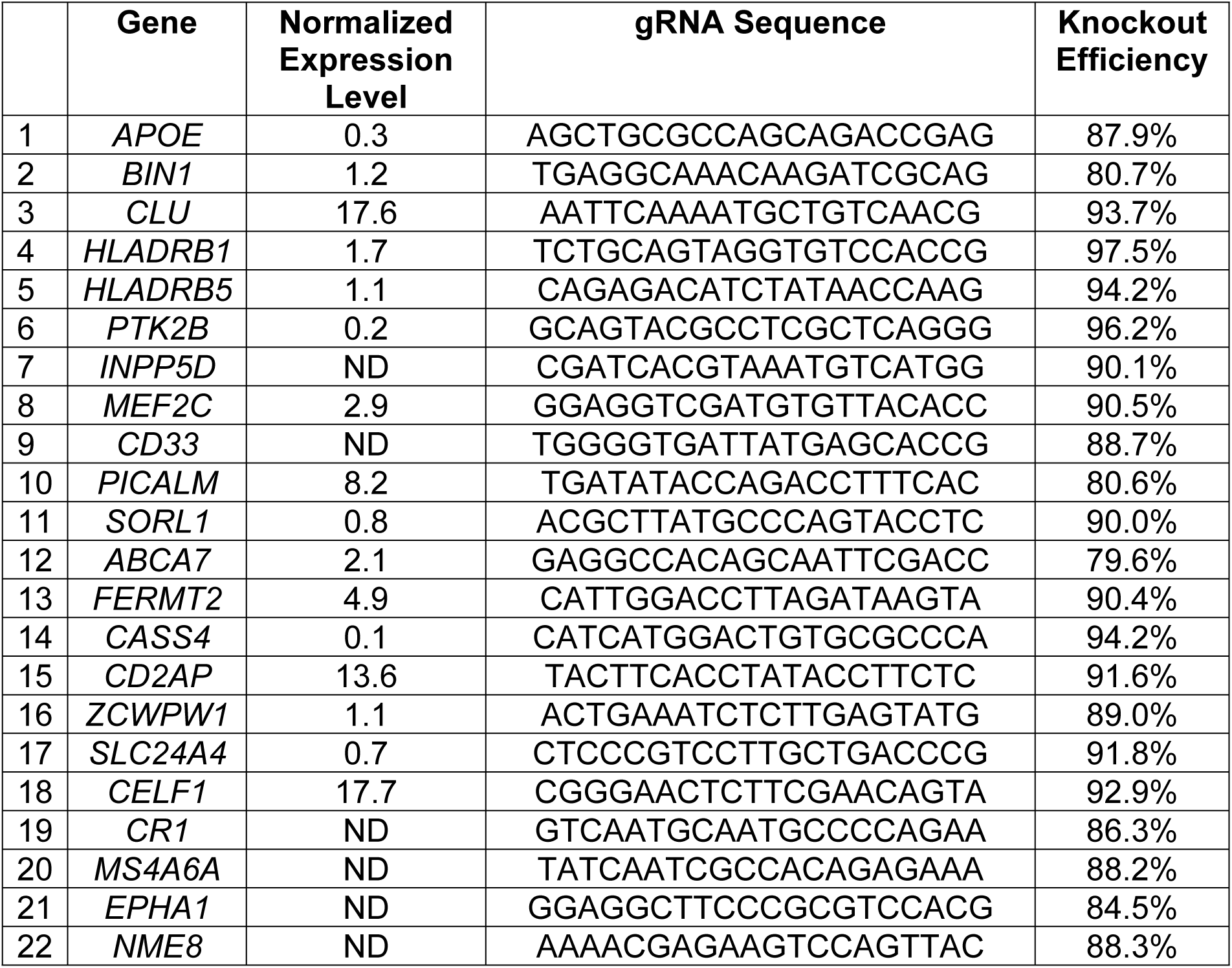
gRNAs used in this study. The knock out efficiency of gRNAs was verified using TIDE. Published expression patterns in HEK293 cells from the <u>Human Protein Atlas</u> are also listed. NX refers to normalized expression levels.

### Uptake assay

HEK293T cells were plated at 15000 cells per well in a 96-well plate. Fluorescently labeled tau aggregates were sonicated (QSonica) for 30s at a setting of 65 (corresponding to ∼80 watts) and were applied to cell media for 4h as per prior studies (18). For positive control in uptake inhibition, fibrils were preincubated overnight at 4°C in media containing heparin at 100 μg/ml. Cells were harvested with 0.05% trypsin and suspended in flow cytometry buffer (HBSS plus 1% FBS and 1 mM EDTA) before quantitation by flow cytometry (LSRFortessa SORP, BD Biosciences). Aggregate internalization was quantified by measuring median fluorescence intensity (MFI) per cell. Technical triplicates were carried out in each condition, and a minimum of 5000 single cells were analyzed per replicate. We determined the average MFI of the replicates for each condition and normalized to aggregate uptake of control sample. Data analysis was performed using FlowJo version 10 software (Treestar, Inc.) and GraphPad Prism version 8 for Windows.

### Seeding assay

A stable monoclonal FRET biosensor cell line overexpressing tau RD(P301S)-C/R was created by selection and amplification of a single cell after viral transduction and culture in puromycin. Biosensor cells were plated at a density of 10,000 cells/well in a 96-well plate. Recombinant tau fibrils were sonicated for 30s at a setting of 65. Aggregates were applied to the cells in volumes of 50μl per well, and incubated for an additional 48h. Tau (50nM) was added directly to the cells after sonication. Alternatively Lipofectamine 2000 (Thermo Fisher Scientific) was used to transduce tau (5nM). After 48h cells were harvested with 0.05% trypsin, fixed in 2% paraformaldehyde for 10 min and then resuspended in flow cytometry buffer (HBSS plus 1% FBS and 1 mM EDTA). We quantified FRET as described previously using the LSRFortessa (29) except that we identified single cells that were both mClover and mRuby positive and subsequently quantified FRET-positive cells within this population. For each data set, three independent experiments with three technical replicates were performed. For each experiment, a minimum of ∼5000 single cells per replicate were analyzed. Data analysis was performed using FlowJo version 10 software and GraphPad Prism version 8.

### Seed maintenance assay

A stable monoclonal LM 39-9 cell line overexpressing tau RD (P301L/V337M) tagged to cyan and yellow fluorescent proteins was used for the tau seed maintenance experiment. These cells had previously been developed for their ability to stably propagate aggregates that enable detection by FRET, as distinct from the first description of the tau RD(P301L/V337M)-YFP cells described previously(7,20). LM 39-9 cells were plated at 10000 cells per well in a 96-well plate. After transduction with virus encoding appropriate gRNA, cells were maintained for 2 weeks prior to analysis. Cells were harvested with 0.05% trypsin and fixed in 2% paraformaldehyde for 10min, and then resuspended in flow cytometry buffer (HBSS plus 1% FBS and 1 mM EDTA). The LSRFortessa SORP (BD Biosciences) was used to perform FRET flow cytometry. FRET was quantified as described previously(23) with the following modification; we identified single cells that were YFP- and CFP-positive and subsequently quantified FRET-positive cells within this population. For each data set, three independent experiments with three technical replicates were performed. For each experiment, a minimum of ∼5000 single cells per replicate were analyzed. Data analysis was performed using FlowJo version 10 software (Treestar Inc.) and GraphPad Prism version 7 for Windows.

## RESULTS

### Knockout of 22 AD GWAS candidates

We first identified the candidate genes based on the reported GWAS of AD (14). Meta-analysis of different GWAS has confirmed the importance of 22 genes as AD risk modifiers (Table 1) (16). We targeted each gene individually with two independent guide RNAs (gRNAs). The gRNAs were cloned into a lentivirus construct (21), and transduced into cultured cells. Cells were cultured for 10 days in the presence of puromycin to select for stable integration of the virus and presumed genetic disruption. We confirmed genetic disruption of each gene using Tracking of Indel by DEcomposition (TIDE), a method based on sequencing the target genes to detect disruption of the sequence through insertion/deletion (indel) at the site of gRNA binding(22). This confirmed high frequency indels at each of the genes targeted by our constructs (see Table 1; Supplementary data-Fig S1 for an example). We selected the gRNAs with high indel efficiency (>80%) in TIDE analysis (Table 1) and used those gRNAs for subsequent assays. We noted that 6 genes are reportedly expressed at very low or undetectable levels in HEK293 cells (Table 1), but carried these through in our analyses nonetheless.

### AD GWAS gene disruption does not affect tau uptake

We have previously determined that heparan sulfate proteoglycans (HSPGs) play a critical role in binding tau aggregates, mediating their uptake and seeding activity (17), (18). Compounds such as heparin or similar small molecules, which bind tau assemblies and compete for their binding to HSPGs, block tau uptake (18). We tested the role of GWAS genes by evaluating HEK293T cells in which we had individually disrupted each. As a positive control, we knocked out NDST1, which we have previously determined to be required for proper HSPG sulfation and to mediate tau uptake (18). We prepared full-length (2N4R) fibrils and labeled them using Alexa Fluor 647 via succinimidyl ester amine reaction. We applied labeled tau fibrils to cultured cells for 4h, followed by washing, trypsin treatment (to digest extracellular tau and release cells from the culture plate), and analysis by flow cytometry according to prior methods (17). We observed no effect of GWAS gene knockout on tau uptake, whereas heparin treatment reduced uptake approximately 90%, and NDST1 knockdown reduced uptake approximately 50% (Fig. 1).

**Figure 1:**
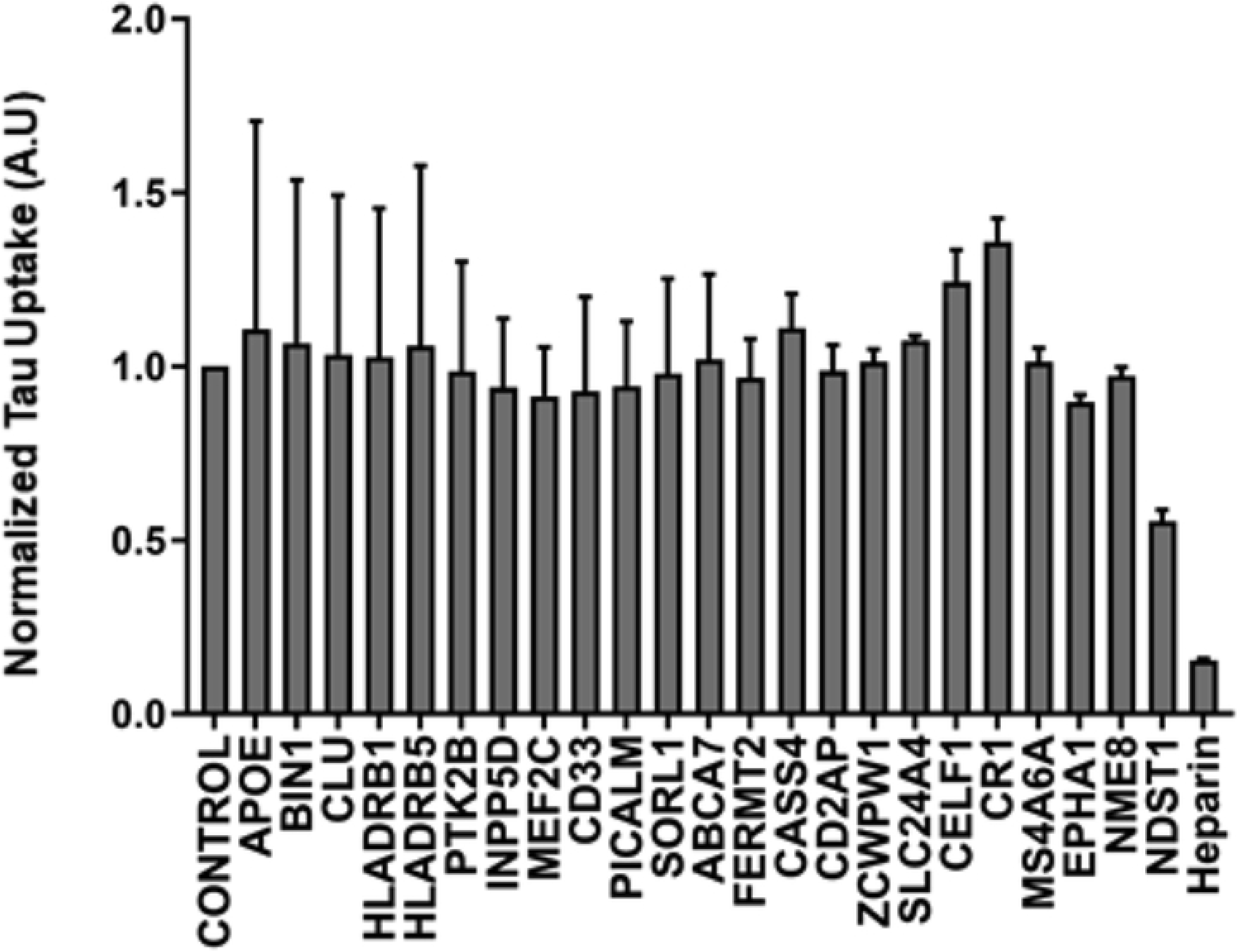
Knockout of GWAS genes does not modify uptake of tau. GWAS Genes were individually targeted in HEK293T cells using CRISPR/Cas9 to create polyclonal knockout cell lines. The cell lines were then tested for internalization of fluorescently labeled tau aggregates by measuring MFI per cell with flow cytometry. None of the gene knockouts changed tau uptake. Data were collected in triplicate and normalized to uptake from control cells treated with scrambled gRNA. The X-axis indicates the targeted genes, and the Y-axis indicates tau uptake relative to scrambled gRNA. Heparin and NDST1 were used as positive controls for uptake inhibition. Error bars indicate the SEM.

### AD GWAS gene disruption does not affect tau seeding

An exogenous tau assembly that gains entry to the cytoplasm acts as a template to convert endogenous tau to a fibrillar form, a process termed “seeding.” Seeding is initiated by application of relatively low concentrations of tau assemblies to the cell media. In the absence of additional reagents, these assemblies bind HSPGs, are internalized, and initiate seeding reactions. This type of seeding, which we have termed “naked,” is relatively inefficient, and typically results in conversion of approximately 1-5% of the cells to an aggregated state. When the tau repeat domain containing a single disease-associated mutation (P301S) is fused to mClover3 (C) or mRuby3 (R) (or a similarly compatible fluorescent protein pair), aggregation enables fluorescence resonance energy transfer (FRET) induced by proximity of the fluorescent proteins. This allows quantitation of seeding activity by flow cytometry.

As an alternative, incubation of tau seeds with Lipofectamine (or a similar reagent) enables transduction of seeds with very high efficiency, approximately 100-fold more than naked seeding. We expressed tau RD(P301S)-C/R in HEK293 cells to form a monoclonal “biosensor” line with high sensitivity to exogenous tau aggregates, similar to a line previously reported (23). To test the role of GWAS genes in the tau seeding process, we knocked out each in the biosensor cells. These were treated with exogenous tau fibrils alone, or with Lipofectamine(19). We measured seeding activity by quantitative flow cytometry. We observed no consistent significant effect of GWAS gene knockout upon naked seeding (Fig. 2) or after Lipofectamine-mediated aggregate delivery (Fig. 3).

**Figure 2.**
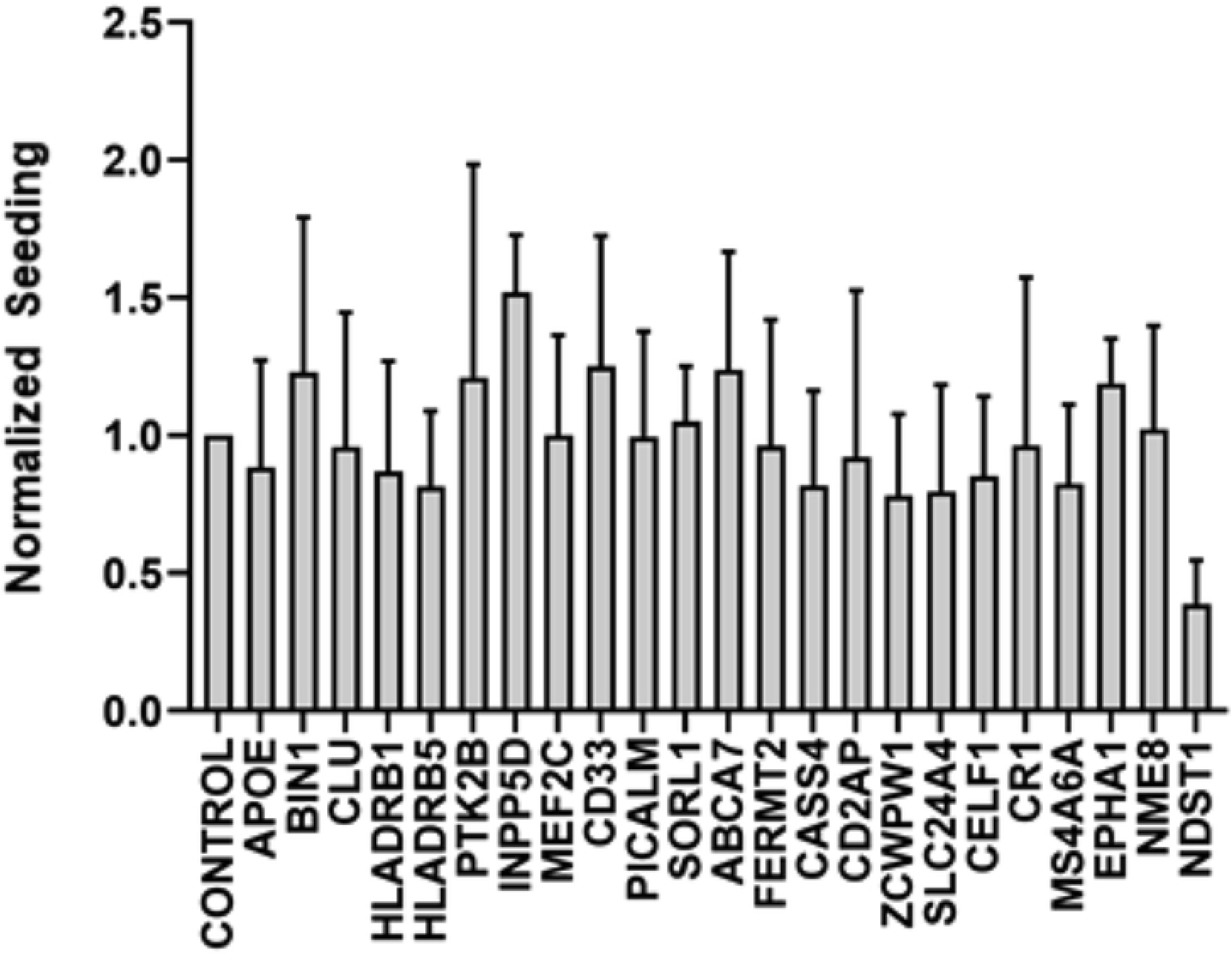
Knockout of AD GWAS genes does not modify naked tau seeding. GWAS Genes were individually targeted in HEK293T RD(P301S)-C/R biosensor cells using CRISPR/Cas9 to create polyclonal knockout cell lines, which were cultured for 2 weeks in the presence of puromycin. Recombinant tau fibrils were added to those cells to induce seeding. Seeding was quantified using FRET, and the percentage of FRET-positive cells was normalized to the scrambled gRNA. Data were collected in triplicate. The X-axis indicates the targeted genes, and the Y-axis indicates normalized seeding activity. None of the genes modified the seeding efficiency. Heparin and NDST1 were positive controls for uptake inhibition. Error bars indicate the SEM.

**Figure 3.**
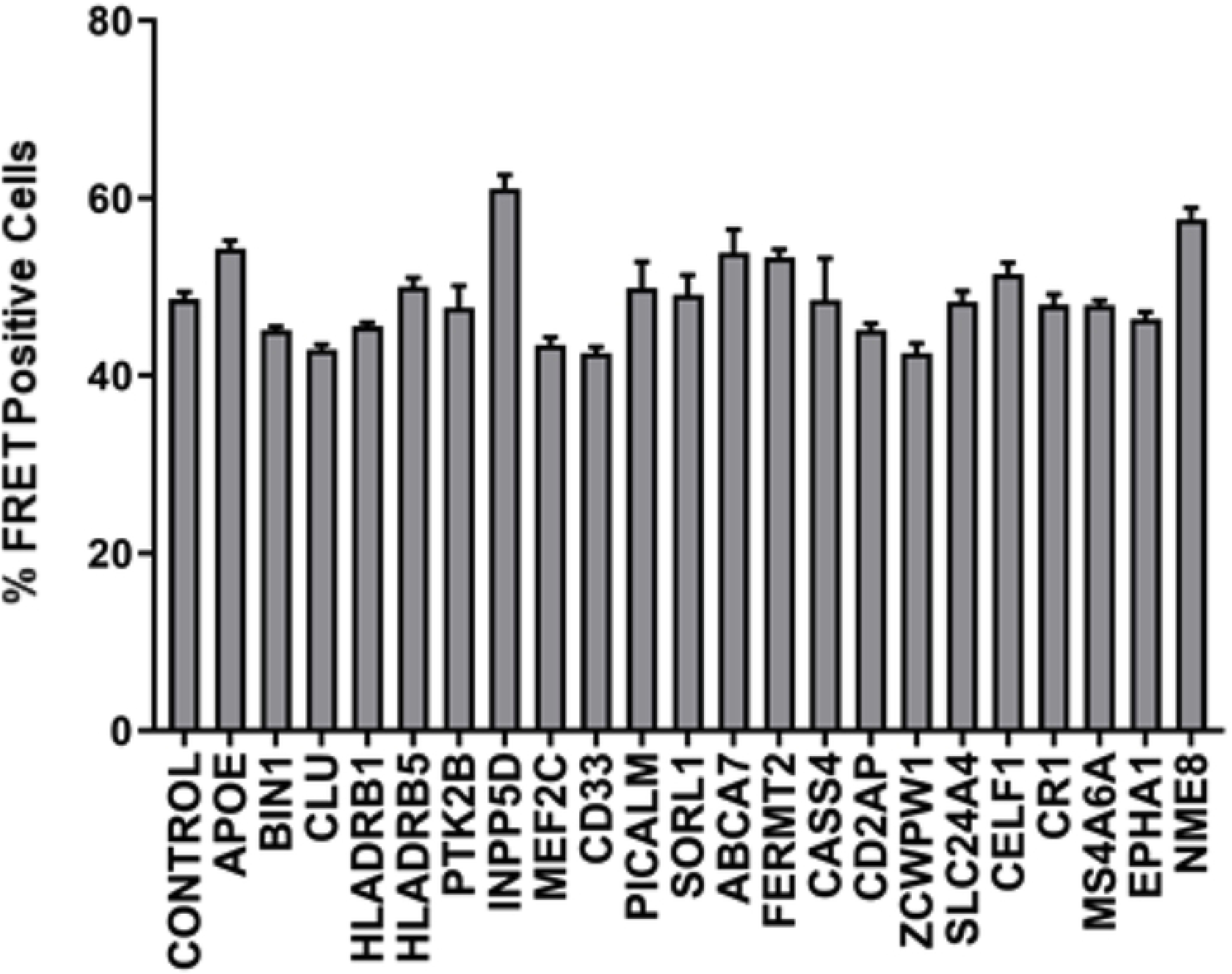
Knockout of AD GWAS genes does not modify Lipofectamine-mediated tau seeding. AD GWAS genes were individually targeted in HEK293T RD(P301S)-C/R biosensor cells using CRISPR/Cas9 to create polyclonal knockout cell lines, which were cultured for 2 weeks in the presence of puromycin. Recombinant tau fibrils were mixed with Lipofectamine to facilitate direct delivery to the cytoplasm. Seeding was quantified using FRET, and the percentage of FRET-positive cells was normalized to the scrambled gRNA. Data were collected in triplicate. The X-axis indicates the targeted genes, and the Y-axis represents normalized seeding activity. No knockout modified seeding efficiency. Error bars indicate the SEM.

### GWAS gene disruption does not affect tau aggregate maintenance

We have previously observed that dividing cells propagate tau aggregates of distinct conformation, termed strains, that transmit pathology between animals, and specify unique pathologies(7,20). Studies of yeast prions indicate that aggregate propagation requires accessory factors, e.g. Hsp104(24), and thus we hypothesized AD GWAS genes might affect this process. We created a cell line that propagated a distinct tau strain, termed LM39-9. These cells constitutively express aggregates of tau RD containing two disease-associated mutations (P301L/V337M) that are fused to cyan and yellow fluorescent proteins (which constitute a FRET pair). LM39-9 cells exhibit high aggregate transmission efficiency (∼99%), which can be easily tracked over time by flow cytometry. We used lentivirus to individually disrupt each of the AD GWAS genes, cultured the LM39-9 cells for 2 weeks, and then quantified the percentage of cells containing aggregates using flow cytometry. We observed no loss of aggregation following disruption of any GWAS gene (Fig. 4), indicating none was critical to aggregate maintenance in this cell model.

**Figure 4.**
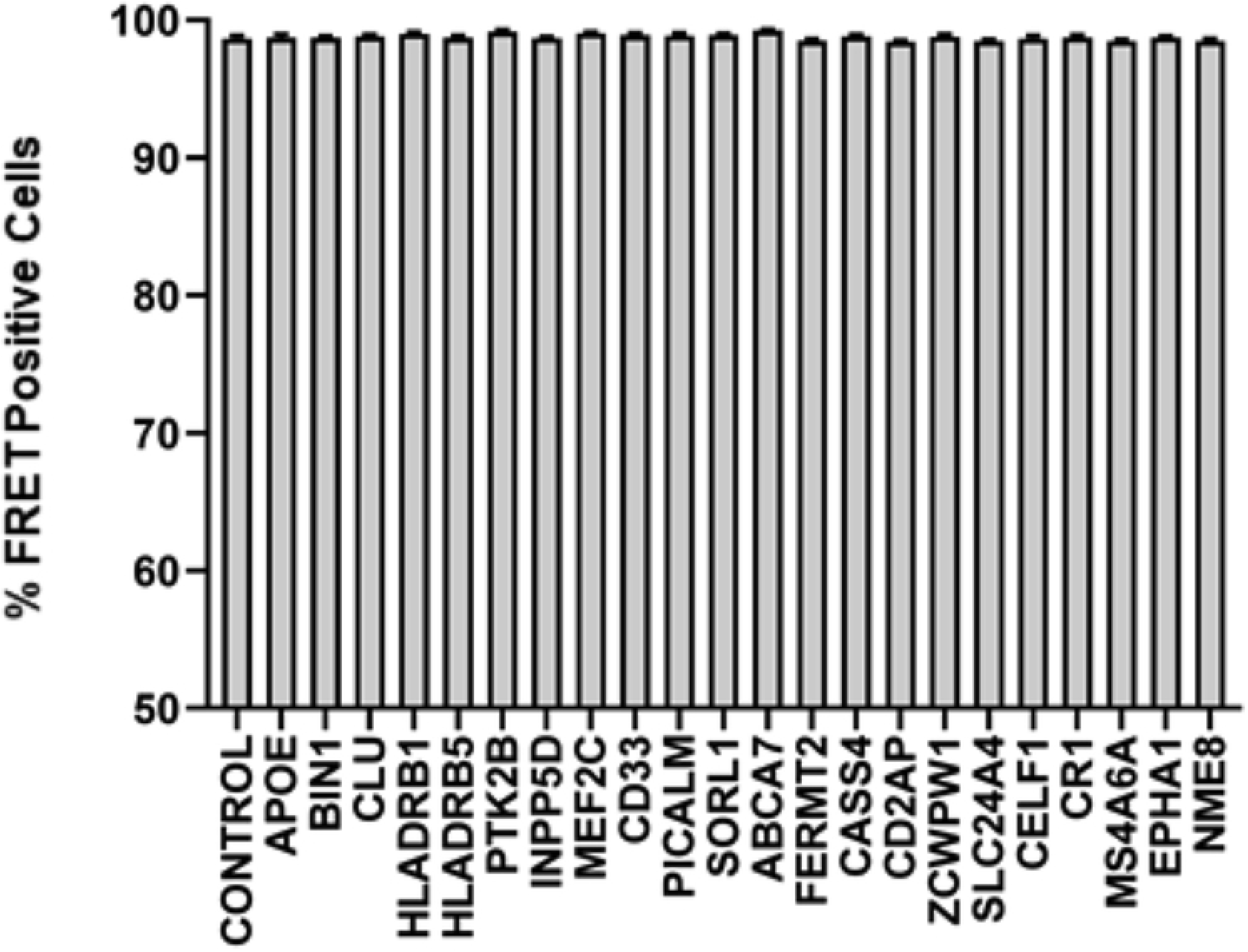
Knock out of AD GWAS genes does not modify tau aggregate maintenance. AD GWAS genes were individually targeted in LM39-9 cells using CRISPR/Cas9 to create polyclonal knockout cell lines, which were cultured for 2 weeks in the presence of puromycin. The cell lines were tested for the loss of tau aggregates using FRET flow cytometry. The X-axis indicates the targeted genes, and the Y-axis represents percentage of FRET-positive cells. None of the genes inhibited aggregate maintenance within the LM-39-9 cell lines. Error bars indicate the SEM.

## DISCUSSION

The mechanisms that govern prion activities of tau are largely unknown. AD GWAS genes are putative modifiers of pathogenesis, and are thus of interest. Consequently in this work we tested the hypothesis that AD GWAS genes would impact uptake, seeding, or aggregate maintenance of tau, which may play a critical role in neurodegeneration. We confirmed each of the 22 gRNAs we studied in fact disrupted their target genes at high frequency. We then studied the effects of the knockouts across a range of putative steps in pathogenesis for which we have previously developed quantitative cell-based assays. We did not observe any impact on knockout in any of the fundamental events of tau aggregate propagation that we can measure in simple cell systems.

AD and related dementias involve progressive accumulation of tau assemblies in neurons and glia. If tau propagation underlies these disorders, it is conceivable that many cellular mechanisms could be specific to cells of the brain. In this case, modeling these processes in simple cultured cell systems as we have done here might not be particularly productive. And, indeed, this may explain why we did not observe any effect of GWAS genes on the fundamental mechanisms of tau pathology we measured in HEK293T cells. 6 of the genes studied (*INPP5D, CD33, CR1, MS4A6A, EPHA1, NME8*) are reportedly not expressed at high levels in these cells, and thus cannot be completely excluded as important for tau prion propagation (although they are clearly not required for this process to occur in HEK293T cells).

Due to the impracticality of optimizing detection methods to measure expression levels of multiple proteins, we confirmed the function of our knockout vectors with sequencing of endogenous genes. Future studies in neurons, which present additional challenges to screening studies we have performed here, would be helpful in this regard. However the HEK293T models have previously proven very useful in defining modes of cell uptake of pathological tau assemblies, seeding, and strain maintenance that have translated well to primary neurons and mouse models (7,17,18). Similarly, these simple systems have readily propagated unique tau strains derived from recombinant fibrils and human tauopathy brains that can be transmitted and propagated in animal models (7,25). We fully recognize that without extension of findings derived from simple systems such as these into animal or even human studies it will be difficult to know how these simple models reflect actual events in the brain.

The relationship of GWAS to AD pathogenesis is complex, as hits may involve genes that are not directly involved in the hypothetically critical process of tau propagation. For example, genes associated with microglial function, such as TREM2, would not be expected to score positive in these studies. Nonetheless, we hope our work will serve as an important reference for those interested in using reductionist cell models to study the role of genes involved in fundamental events of tau propagation.

## ACKNOWLEDGEMENTS

The study was supported by the Rainwater Charitable Foundation and the Crowley Foundation.

---

**Supplementary Figure 1: Analysis of gene editing efficiency by Tracking of Indels by DEcomposition (TIDE).**

(**A**) Representative image of one of the genes targeted with CRISPR/Cas9. After lentivirus exposure, cells were cultured for 2 weeks in puromycin. DNA was isolated from control (scrambled gRNA treated) and *CASS4* KO, and Sanger sequenced. The gRNA sequence used for making the knockout is shown in bold. The predicted Cas 9 cut site is shown as a red dotted line. The base pair number is labelled. The insertions/deletions (N) can be seen in the *CASS4* KO. (**B**) Representative image from TIDE analysis of *CASS4*. Plot shows the overlay of sequence between scrambled control and *CASS4*. The increase in aberrant sequence after the expected cut site is evident in the *CASS4* KO sample, indicating effective gene disruption. (**C**) Plots represent the spectrum of indels and their frequencies for *CASS4*. R^2^ = 0.94. The plot was analyzed and derived from the TIDE web tool (https://tide.deskgen.com/).

